# On the use of calcium deconvolution algorithms in practical contexts

**DOI:** 10.1101/871137

**Authors:** Mathew H. Evans, Rasmus S. Petersen, Mark D. Humphries

## Abstract

Calcium imaging is a powerful tool for capturing the simultaneous activity of large populations of neurons. Here we determine the extent to which our inferences of neural population activity, correlations, and coding depend on our choice of whether and how we deconvolve the calcium time-series into spike-driven events. To this end, we use a range of deconvolution algorithms to create nine versions of the same calcium imaging data obtained from barrel cortex during a pole-detection task. Seeking suitable values for the deconvolution algorithms’ parameters, we optimise them against ground-truth data, and find those parameters both vary by up to two orders of magnitude between neurons and are sensitive to small changes in their values. Applied to the barrel cortex data, we show that a substantial fraction of the processing methods fail to recover simple features of population activity in barrel cortex already established by electrophysiological recordings. Raw calcium time-series contain an order of magnitude more neurons tuned to features of the pole task; yet there is also qualitative disagreement between deconvolution methods on which neurons are tuned to the task. Finally, we show that raw and processed calcium time-series qualitatively disagree on the structure of correlations within the population and the dimensionality of its joint activity. Collectively, our results show that properties of neural activity, correlations, and coding inferred from calcium imaging are sensitive to the choice of if and how spike-evoked events are recovered. We suggest that quantitative results obtained from population calcium-imaging be verified across multiple processed forms of the calcium time-series.

## 1 Introduction

Calcium imaging is a wonderful tool for high yield recordings of large neural populations (Harris et al., 2016; Stringer et al., 2019a; Ahrens et al., 2013; Portugues et al., 2014). Many pipelines are available for moving from pixel intensity across frames of video to a time-series of calcium fluorescence in the soma of identified neurons (Mukamel et al., 2009; Vogelstein et al., 2010; Kaifosh et al., 2014; Pachitariu et al., 2016; Deneux et al., 2016; Pnevmatikakis et al., 2016; Friedrich et al., 2017; Keemink et al., 2018; Giovannucci et al., 2019). As somatic calcium is proportional to the release of spikes, so we wish to use these fluorescence time-series as a proxy for spiking activity in large, identified populations of neurons. But raw calcium fluorescence is slow on the time-scale of spikes, nonlinearly related to spiking, and contains noise from a range of sources.

These issues have inspired a wide range of deconvolution algorithms (Theis et al., 2016; Berens et al., 2018; Stringer and Pachitariu, 2018), which attempt to turn raw somatic calcium into something more closely approximating spikes. Deconvolution algorithms themselves range in complexity from simple deconvolution with a fixed kernel of the calcium response (Yaksi and Friedrich, 2006), through detecting spike-evoked calcium events (Jewell and Witten, 2018; Pachitariu et al., 2016), to directly inferring spike times (Vogelstein et al., 2010; Lütcke et al., 2013; Deneux et al., 2016). This continuum of options raises the further question of the extent to which we should process the raw calcium signals. We address here the question facing any systems neuroscientist using calcium imaging: do we use the raw calcium, or attempt to clean it up? Thus our aim is to understand if our choice matters: to what extent do our inferences about neural activity, correlations, and coding depend on our choice of raw or deconvolved calcium time-series.

We proceed here in two stages. In order to use deconvolution algorithms, the data analyst needs to choose their parameters. We thus first address how good these algorithms can be in principle with optimised parameters, and how sensitive their results are to the choice of parameter values. To do so, we evaluate qualitatively different deconvolution algorithms by optimising their parameters against ground truth data with known spikes.

With our understanding of their parameters in hand, we then turn to our main question, by analysing a large-scale population recording from the barrel cortex of a mouse performing a whisker-based decision task. We compare estimates of population coding and correlations obtained using either raw calcium signals, or a range of time-series derived from those calcium signals, covering simple deconvolution, event detection, and spikes.

We find that a substantial fraction of the deconvolution methods used here fail to recover basic features of population activity in barrel cortex established from electrophysiology. The inferences we draw about coding qualitatively differ between raw and deconvolved calcium signals. In particular, coding analyses based on raw calcium signals detect an order of magnitude more neurons tuned to task features. Yet there is also qualitative disagreement between deconvolution methods on which neurons are tuned. The inferences we draw about correlations between neurons do not distinguish between raw and deconvolved calcium signals, but can qualitatively differ between deconvolution methods. Our results thus suggest care is needed in drawing inferences from population recordings of somatic calcium, and that one solution is to replicate all results in both raw and deconvolved calcium signals.

## 2 Results

### 2.1 Performance of deconvolution algorithms on ground-truth data-sets

We select here three deconvolution algorithms that infer discrete spike-like events, each an example of the state of the art in qualitatively different approaches to the problem: Suite2p (Pachitariu et al., 2016), a peeling algorithm that matches a scalable kernel to the calcium signal to detect spike-triggered calcium events; LZero (Jewell and Witten, 2018), a change-point detection algorithm, which finds as events the step-like changes in the calcium signal that imply spikes; and MLspike (Deneux et al., 2016), a forward model, which fits an explicit model of the spike-to-calcium dynamics in order to find spike-evoked changes in the calcium signal, and returns spike times. We emphasis that these methods were chosen as exemplars of their approaches, and are each innovative takes on the problem; we are not here critiquing individual methods, nor are we seeking a “best” method. Rather, we are using an array of methods to illustrate the problems and decisions facing the data analyst when using calcium imaging data.

We first ask if these deconvolution methods work well in principle, by testing if there exists parameter sets for which they each successfully recover known spike times from calcium traces. We fit the parameters of each method to a data-set of 21 ground-truth recordings (Chen et al., 2013), where the spiking activity of a neuron is recorded simultaneously with a high-signal-to-noise cell-attached glass pipette and 60 Hz calcium imaging (Figure 1a). To fit the parameters for each recording, we sweep each method’s parameter space to find the parameter value(s) with the best match between the true and inferred spike train.

**Figure 1:**
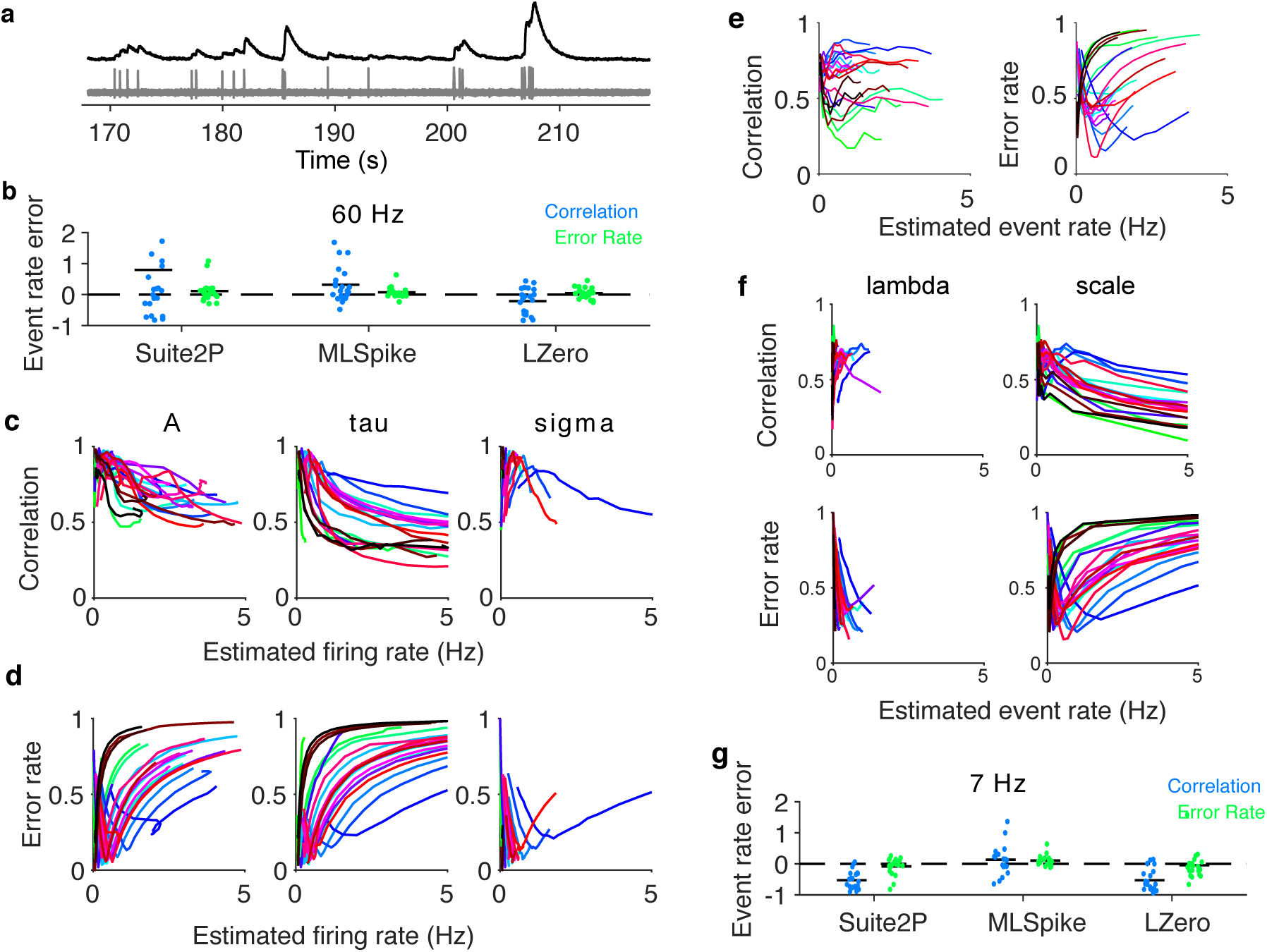
Deconvolution algorithms can accurately recover spiking events in principle. (a) Example simultaneous recording of somatic voltage (grey) and calcium activity (black) imaged at 60Hz. (b) Error in estimating the true firing rate when using optimised parameters, across all three methods. One symbol per recording. We separately plot errors for parameters optimised to maximise the correlation coefficient, and the errors for parameters optimised to minimise the error rate. Horizontal black bars are means. Error is computed relative to the true firing rate: (*Rate*_*true*_ − *Rate*_*estimated*_*/Rate*_*true*_); and error of 1 thus corresponds to twice as many estimated spikes as there are in the ground-truth data. For LZero and Suite2p, *Rate*_*estimated*_ is computed from event times. (c) Dependence of MLspike’s deconvolution performance on the firing rate of the inferred spike train. For each of MLSpike’s free parameters, we plot the correlation coefficient between true and inferred spikes as a function of the firing rate estimated from the inferred spikes obtained at each tested parameter value. One line per recording; colours are used solely to help distinguish the lines. Parameters: *A*, calcium transient amplitude per spike (Δ*F/F*); *τ*, calcium decay time constant (s); *σ*, background (photonic) noise level (Δ*F/F*) (d) as in (c), but using error rate between the true and inferred spikes. (e) Dependence of Suite2p’s deconvolution performance on the firing rate of the inferred event train as its detection threshold parameter is varied. Left: correlation coefficient; right: error rate. (f) Dependence of LZero’s deconvolution performance on the firing rate of the inferred event train, as its two parameters are varied: *λ*, sparsity of spike events; scale, the magnitude of a single spike-induced fluorescence change. (g) As for (b), but with the somatic calcium down-sampled to 7Hz before optimising parameters for the deconvolution methods.

The best-fit parameters depend strongly on how we evaluate the match between true and inferred spike trains. The Pearson correlation coefficient between the true and inferred spike train is a common choice (Brown et al., 2004; Paiva et al., 2010; Theis et al., 2016; Reynolds et al., 2018; Berens et al., 2018), typically with both trains convolved with a Gaussian kernel to allow for timing errors. However, we find that choosing parameters to maximise the correlation coefficient can create notable errors. The inferred spike trains from MLSpike have too many spikes on average (mean error over recordings: 31.72%), and the accuracy of recovered firing rates widely varies across recordings (Fig 1b, blue symbols). We attribute these errors to the noisy relationship between the correlation coefficient and the number of inferred spikes (Figure 1c): for many recordings, there is no well-defined maximum coefficient, especially for the amplitude parameter *A*, so that near-maximum correlation between true and inferred trains is consistent with a wide range of spike counts in the inferred trains. We see the same sensitivity for the event rates from recordings optimised using Suite2p (Figure 1e) and LZero (Figure 1f, top). If we compare their inferred event rates to true firing rates (Fig 1b), we see Suite2p estimates far more events than spikes (mean error 79.47%) and LZero fewer events than spikes (mean error: - 21.14%). These further errors are problematic: there cannot be more spike-driven calcium events than spikes, and LZero’s underestimate is considerably larger than the fraction of frames with two or more spikes (*<* 0.002% frames).

To address the weaknesses of the Pearson correlation coefficient, we instead optimise parameters using the error rate metric of Deneux et al. (2016). The error rate is derived from the proportions of missed and excess spikes (see Methods), and returns a normalised score between 0 for a perfect match between two spike trains, and 1 when all the spikes are missed. This comparison between inferred and true spike trains is most straightforward for algorithms like MLSpike that directly return spike times; for the other algorithms, we use here their event times as inferred spikes, a reasonable choice given the low firing rate and well separated spikes in the ground truth data. Choosing parameters to minimise the error rate between the true and inferred spike-trains results in excellent recovery of the true number of spikes for all three deconvolution methods (Fig 1b, green symbols), with mean errors in spike counts of 12% excess spikes for Suite2P, 7.3% for MLSpike, and 5% for LZero. As we show in Figure 1d-f, for all three deconvolution methods the error rate has a well-defined minimum for almost every recording. Consequently, all deconvolution methods can, in principle, accurately recover the true spike-trains given an appropriate choice of parameters.

A potential caveat here is that the ground-truth data are single neurons imaged at a frame-rate of 60Hz, an order of magnitude greater than is typically achievable in population recordings (Peron et al., 2015a). Such a high frame-rate could allow for more accurate recovery of spikes than is possible in population recordings. To test this, we downsample the ground-truth data to a 7Hz frame-rate, and repeat the parameter sweeps for each deconvolution method applied to each recording. As we show in Figure 1g, optimising parameters using the minimum error rate still results in excellent recovery of the true spike rate (and interestingly for some recordings reduces the error when using the correlation coefficient). Lower frame-rates need not then be an impediment to using deconvolution methods.

### 2.2 Parameters optimised on ground-truth are widely distributed and sensitive

What might be an impediment to using deconvolution methods on population recordings is if the best parameter values vary widely between neurons. If so, then parameters optimised for one neuron would generalise poorly to the rest of the population.

Figure 2a-b plots the best-fit parameter values for each single neuron recording across deconvolution methods and sampling rates. Each method has at least one parameter with substantial variability across recordings, varying by an order of magnitude or more. This suggests that the best parameters for one neuron may not apply to another. In turn, this parameter variation between neurons could mean that analysis of population recordings created from a single set of deconvolution parameters would potentially include many aberrant time-series.

**Figure 2:**
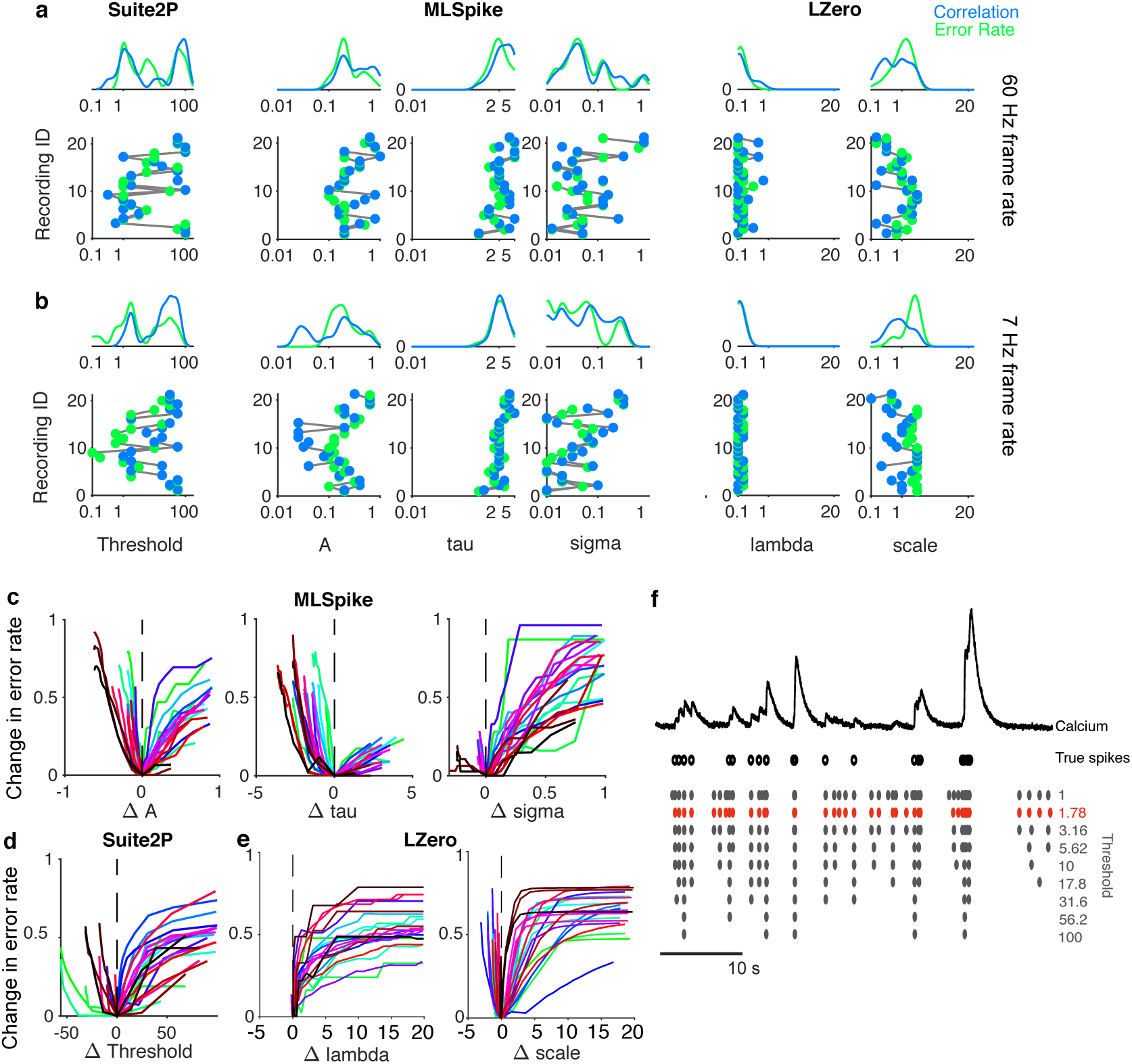
Variation in best-fit spike deconvolution parameters across ground-truth recordings. (a) Distributions of optimised parameter values across recordings. For each parameter (a column), the bottom panel plots the found parameter values on the x-axis against the recording ID on the y-axis (in an arbitrary but consistent order); the top panel plots the marginal distribution of the parameter value over all recordings. We plot for each recording the optimised parameter value found using correlation coefficient and error rate. Lines join recordings from the same neuron. (b) As for panel (a), fits to the same ground-truth data down-sampled to 7 Hz. (c) Change in error rate as a function of the change away from a parameter’s optimum value, for each of MLSpike’s free parameters. One line per recording, at 60 Hz frame rate. (d) As for panel (c), for changes in Suite2p’s threshold parameter. (e) As for panel (c), for changes in LZero’s two parameters. (f) Example of the range of inferred spike event trains possible when applying plausible but wrong parameter values to a recording. For one recording, we plot in red the inferred spike events detected using its optimised threshold parameter for Suite2p. Alongside we plot the inferred trains of spike events that result if we vary the threshold parameter across the range of optimised values found within the set of 21 recordings (values in panel (a), optimised using error rate).

The problem of between-neuron variation in parameter values would be compensated somewhat if the quality of the inferred spike or event trains is robust to changes in those values. However, we find performance is highly sensitive to changes in some parameters. Figure 2c-e shows that for most recordings the quality of the inferred spike train abruptly worsens with small increases or decreases in the best parameter, regardless of the deconvolution method used. As we show in Figure 2f, the inferred spikes for a single neuron can vary dramatically as we change a parameter value, even when we restrict ourselves to just the range of optimised values across the recordings. That the parameters are sensitive and vary considerably across neurons has the significant implication that, unless ground truth data is available for every neuron being analysed, deconvolution algorithms could be substantially inaccurate.

### 2.3 Deconvolution of population imaging in barrel cortex during a decision task

We turn now to the core problem facing any analyst of population calcium imaging data: there are rarely ground truth data, and never for every neuron. In the absence of ground-truth data, there is no way of selecting a “best” deconvolution algorithm or a “best” set of parameters for analysing a population recording. Yet the above results imply that the insights we gain about population activity would indeed depend crucially on which deconvolution method we use. We now test the extent of this dependence by applying 8 different deconvolution methods to the same raw calcium time-series, and compare the resulting statistics of neural activity, properties of neural coding, and the extent and structure of correlations between neurons.

The data we use are two-photon calcium imaging time-series from a head-fixed mouse performing a whisker-based two-alternative decision task (Fig. 3a-b), from the study of Peron et al. (2015b). We analyse here a single session with 1552 simultaneously recorded pyramidal neurons in L2/3 of a single barrel in somatosensory cortex, imaged at 7 Hz for just over 56 minutes, giving 23559 frames in total across 335 trials of the task.

**Figure 3:**
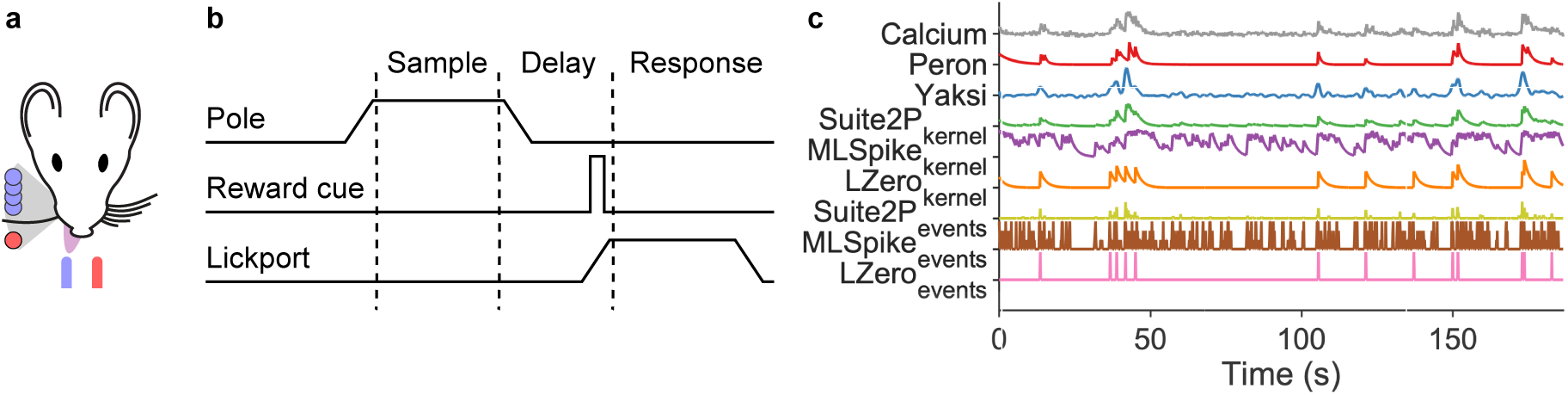
Experimental data from Peron et al. (2015b). (a) Schematic of task set-up. A pole was raised within range of the single right-side whisker; the pole’s position, forward (red circle) or backward (blue circles) indicated whether reward would be available from the left or right lick-port. (b) Schematic of trial events. The pole was raised and lowered during the sample period; a auditory cue indicated the start of the response period. (c) All deconvolution methods applied to one raw calcium signal from the same neuron.

Our primary goal is to understand how the choices of deconvolving these calcium-imaging data alter the scientific inferences we can draw. As our baseline, we use the “raw” Δ*F/F* time-series of changes in calcium indicator fluorescence. We use the above three discrete deconvolution methods to extract spike counts (MLSpike), event occurrence (LZero), or event magnitude (Suite2p) per frame. Given the above-demonstrated dependence of these algorithms on their parameters, we use Yaksi and Friedrich (2006)’s simple deconvolution of the raw calcium with a fixed kernel of the GCaMP6s response to a single spike, whose only free parameters are fixed from data. For comparison, we use Peron et al. (2015b)’s own version of denoised calcium time-series, which they created using a custom version of the peeling algorithm (Lütcke et al., 2013), a greedy template-fitting event-detection algorithm with variable rise and decay time constants across events. The Peron time-series are then the detected spike-events convolved with a kernel of the detected rise and decay time. And finally, for comparison with the Peron time-series, we create equivalent versions for our three discrete-deconvolution methods, by convolving their recovered spikes/events with a fixed GCaMP6s spike-response kernel. Figure 3c show an example raw calcium time-series for one neuron, and the result of applying each of these 8 processing methods. We thus repeat all analyses on 9 different sets of time-series extracted from the same population recording.

We choose the algorithm parameters as follows. Simple deconvolution (Yaksi and Friedrich, 2006) uses a parameterised kernel of the GCaMP6s response to a single spike. For the three discrete deconvolution methods, we choose the modal values of the best-fit parameters that optimised the error rate over the ground-truth recordings. This seems a reasonably consistent way obtaining comparable results between methods, by using the most consistently performing values obtained from comparable data: neurons in the same layer (L2/3) in the same species (mouse), in another primary sensory area (V1). Most importantly for our purposes, choosing the modal values means we avoid extreme and potentially pathological regions of the parameter space. Again this recapitulates the problem facing any analyst of population calcium imaging data, of how to choose the parameters for a deconvolution algorithm in the absence of any ground-truth recordings.

### 2.4 Deconvolution methods disagree on estimates of simple neural statistics

We first check how well each approach recovers the basic statistics of neural activity event rates in L2/3 of barrel cortex. Electrophysiological recordings have shown that the distribution of firing rates across neurons in a population is consistently long-tailed, and often log-normal, all across rodent cortex (Wohrer et al., 2013). Cell-attached recordings of L2/3 neurons in barrel cortex are no different (O’Connor et al., 2010), with median firing rates less than 1 Hz, and a long right-hand tail of rarer high-firing neurons. We thus test if the calcium event rates or spike rates from our time-series follow such a distribution. (Event rates for raw calcium, Peron, Yaksi and the continuous (kernel) versions of the data was obtained by thresholding the calcium time-series)

Figure 4a shows that the raw calcium and two of the discrete deconvolution methods (Suite2p_*events*_, LZero_*events*_) qualitatively match the expected distributions of event rates (median near zero, long right-hand tails). The Peron time-series also have the correct distribution of event rates. All other methods give wrong distributions, whether of spike rates (MLSpike) or event rates (all other methods). There is also little overlap in the distributions of spike rates between the three discrete deconvolution methods. Applying a kernel to their inferred spikes/events shifts rather than smooths the firing rate distributions (Suite2P_*kernel*_, MLSpike_*kernel*_, LZero_*kernel*_), suggesting noise in the deconvolution process is amplified through the additional steps of convolving with a kernel and thresholding.

**Figure 4:**
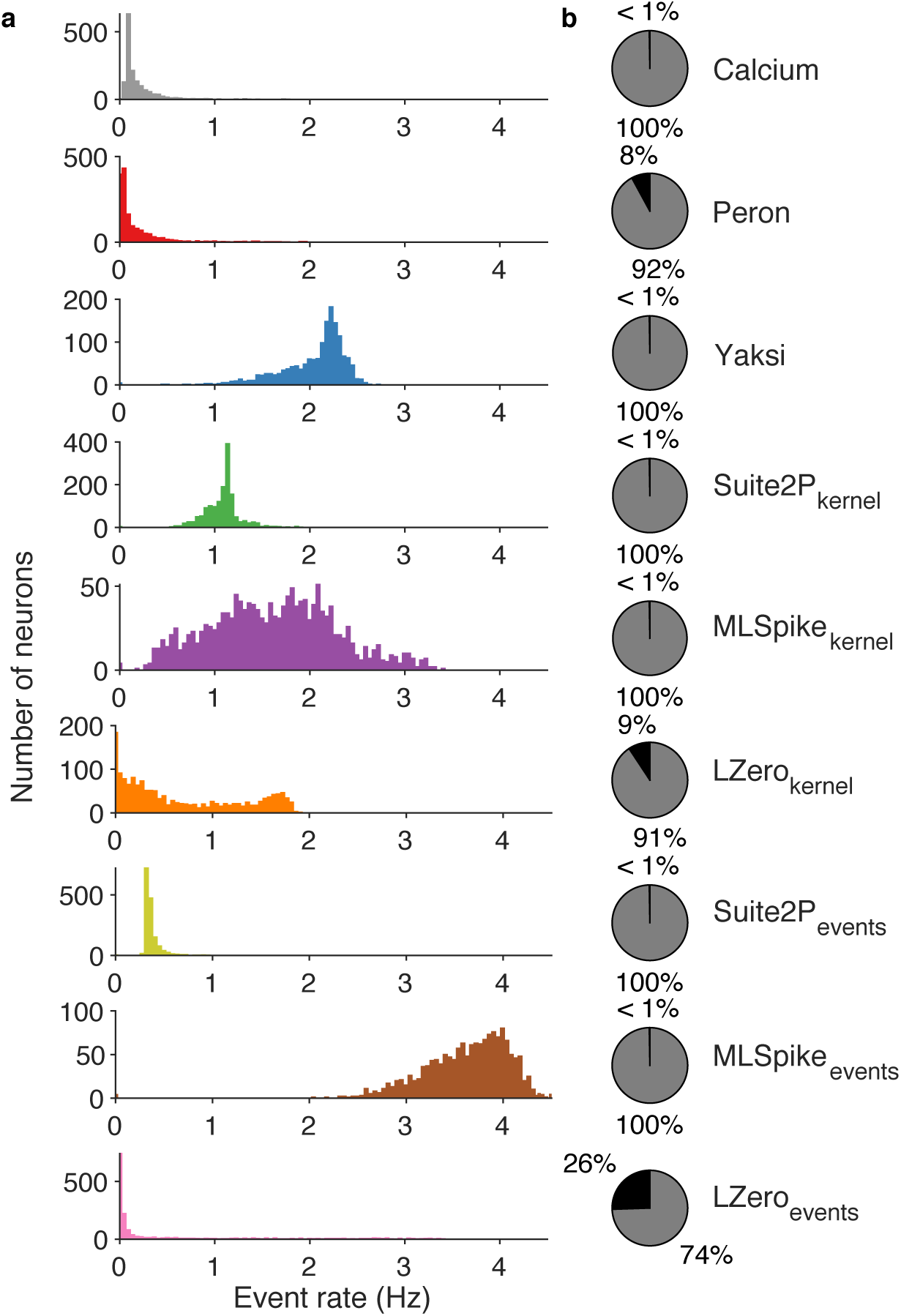
Estimates of population-wide event rates vary qualitatively across deconvolution methods. (a) The distribution of event rate per neuron across the recorded population, according to each deconvolution method. For raw calcium and the five continuous versions of the time-series (upper 6 panels), events are detected as fluorescence transients greater in magnitude than three standard deviations of background noise. The discrete deconvolution methods (lower 3 panels) return per frame: a spike count (MLSpike), a binary event detection (LZero), or an event magnitude (Suite2p); these time-series were thus sparse, with most frames empty. (b) Proportion of active (gray) and silent (black) neurons for each method. Silent neurons are defined following (Peron et al., 2015b) as those with an event rate less than 0.0083Hz.

Cell-attached recordings in barrel cortex have shown that *∼*26% of L2/3 pyramidal neurons are silent during a similar pole localisation task, with silence defined as emitting fewer than one spike every two minutes (O’Connor et al., 2010). For the nine approaches we test here, six estimated the proportion of silent neurons to be less than 1%, including two of the discrete deconvolution methods (Figure 4b). For raw calcium and methods returning continuous time-series, raising the threshold for defining events will lead to more silent neurons, but at the cost of further shifting the event rate distributions towards zero. Even for simple firing statistics of neural activity, the choice of deconvolution method gives widely differing, and sometimes wrong, results.

### 2.5 Inferences of single neuron tuning differ widely between raw calcium and deconvolved methods

In any paradigm where one records the responses of neurons as an animal performs some task, a basic question is what fraction of neurons in a target brain region are selective to some aspect of the task. Here we ask how the detection of task-tuned neurons depends on our choice of processing method for the raw calcium time-series.

The decision task facing the mouse (Fig. 3a) requires that it moves its whisker back- and-forth to detect the position of the pole, delay for a second after the pole is withdrawn, and then make a choice of the left or right lick-port based on the pole’s position (Fig. 3b). As the imaged barrel corresponds to the single spared whisker (on the contralateral side of the face), so the captured population activity during each trial likely contains neurons tuned to different aspects of the task.

Following Peron et al. 2015a, we define a task-tuned neuron as one for which the peak in its trial-averaged activity exceeds the predicted upper limit from shuffled data (Fig. 5a). When we apply this definition to the raw calcium time-series, close to half the neurons are tuned (734/1552; Fig.5b). This is more than double the proportion of tuned neurons we find for the next nearest method (Yaksi’s simple deconvolution), and at least a factor of 5 greater than the proportion of tuned neurons resulting from any discrete deconvolution method (“events”), which each report less than 10% of the neurons are tuned.

**Figure 5:**
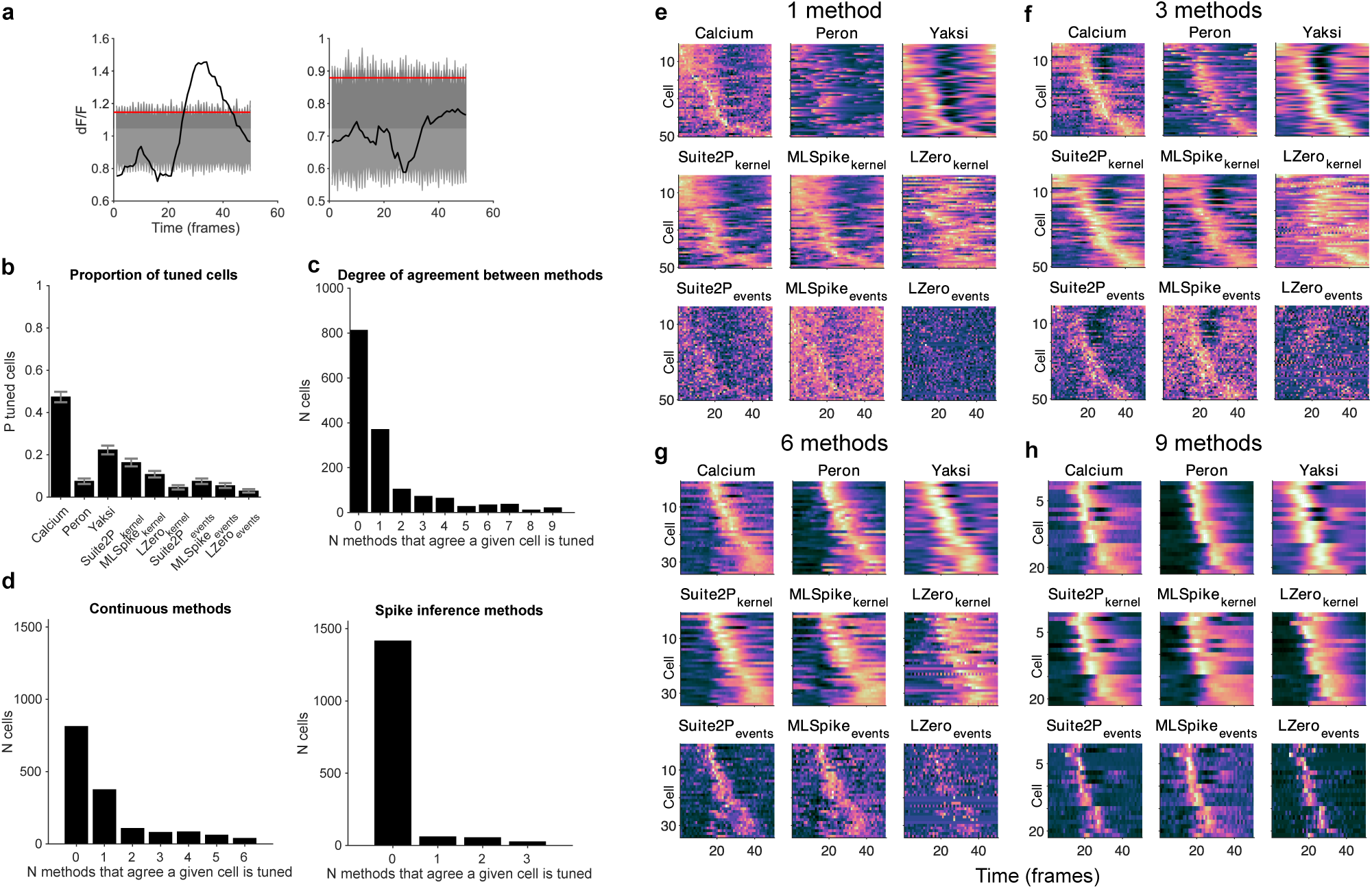
Inferences of single neuron tuning show poor agreement between raw calcium and deconvolution methods, and between methods. (a) Examples of a tuned (left) and non-tuned (right) neuron from the raw calcium time-series. Black: trial-averaged calcium fluorescence. Grey shading: full range of ΔF/F from the shuffled data. Red line: 95th percentile of the peak ΔF/F value across the shuffled data. (b) Number of tuned neurons per deconvolution method. Error bars are 95% Jeffreys confidence intervals for binomial data (Brown et al., 2001). (c) Agreement between methods. For each neuron, we count the number of methods (including raw calcium) for which it is labelled as tuned. Bars show the number of neurons classified as tuned by exactly *N* methods. (d) Similar to (c), but breaking down the neurons into: agreement between methods (including raw calcium) resulting in continuous signals (left panel); and agreement between discrete deconvolution methods (right panel). (e-h) Identifying robust neuron tuning. Panel groups (e) to (h) show neurons classed as tuned by increasing numbers of deconvolution methods. Each panel within a group plots one neuron’s normalised (z-scored) trial-average histogram per row, ordered by the time of peak activity. The first panel in a group of 9 shows histograms from raw calcium signals; each of the 8 subsequent panels shows trial-average histograms for the same neurons, but following processing by each of the eight deconvolution methods.

The wide variation in numbers reflects little consistent agreement between the nine sets of time-series about which neurons are tuned. A substantial fraction of the neurons are found tuned in only one time-series of the nine (Fig.5c). And that time-series is overwhelmingly the raw calcium: of the 734 tuned neurons in the raw calcium time-series, half (364, 49.5%) are unique, detected only in those time-series. By contrast, across all 8 deconvolution methods only 6 neurons are found tuned by one method alone. Thus either the raw calcium time-series contains many erroneously-detected tuned neurons, or the 8 deconvolution methods combined miss many tuned neurons, or both.

One likely source of this broad disagreement is that the raw calcium time-series allows a generous definition of “tuned”. Spike-evoked changes to the somatic calcium concentration are slow on the time-scale of spikes, and the calcium sensors are slower still: the GCaMP6s sensor’s response to a single spike has a rise-to-maximum time of around 0.2 seconds, and a half-width decay time of at least 0.5 seconds (Chen et al., 2013; Dana et al., 2014). This creates a strong low-pass filtering of the underlying spike train, leading to weak correlation between the timing of the spikes and the timing of the calcium changes (see also Sabatini (2019)). Consequently, in the raw calcium time-series a neuron could emit a spike on each trial that are over a second apart between trials, yet each would still contribute to a “peak” in the trial-averaged signal. (Indeed in Fig.5b we see this low-pass filtering effect in the convolved versions of our discrete-deconvolution time-series: we always get more tuned neurons in the “kernel” versions despite them having identical underlying spike-events to the “event” versions.) Finding a tuned neuron in the raw calcium time-series tells us only that the neuron was active during the trial, not that its spikes were specifically tuned to some event in the world.

Indeed, the need to correct the low correlation between raw calcium time-series and behaviour or events in the world is a major reason why deconvolution methods have been developed. Simply convolving the raw calcium signal with a fixed-parameter kernel as in the Yaksi method immediately halves the number of apparently tuned neurons, potentially because the immediate rise time and fixed decay time of the kernel reduce the low-pass filtering of the spike train. And when recovering discrete spike-events a neuron can only be “tuned” if those spike-events align in time across trials, leading to far fewer tuned neurons. We can see then our choice of time-series processing creates a continuum of definitions of neuron tuning in this analysis.

But the implicit definition of tuning is not the only source of disagreement between the 9 time-series. Of the neurons found tuned in more than one time-series, the agreement is still poor. Just 21 (1.35%) are labelled as tuned in all nine (Fig.5c). Even separately considering the continuous and spike-event time-series, we find only 38 (2.4%) neurons are tuned across all six continuous methods, and 25 (1.6%) neurons for all three spike-event deconvolution methods (Fig.5d). Even between time-series with similar implicit definitions of “tuned”, there is inconsistency about which neurons fit that definition.

An approach for the consistent detection of tuned neurons is to find those agreed between the raw calcium time-series and more than one deconvolution method. In Figure 5e-h, we show how increasing the number of methods required to agree on a neuron’s tuned status creates clear agreement between time-series processed with all methods, even if a particular method did not reach significance for that neuron. Even requiring agreement between the raw calcium and just two other methods is enough to see tuning of many neurons. More reliable identification of task-tuned neurons could potentially be achieved by triangulating the raw calcium with the output of multiple deconvolution methods.

In the pole detection task considered here, neurons tuned to pole contact are potentially crucial to understanding the sensory information used to make a decision. Touch onset is known to drive a subset of neurons in barrel cortex to spike with short latency and low jitter (O’Connor et al., 2010; Hires et al., 2015). Detecting such rapid, precise responses in the slow kinetics of calcium imaging is challenging, suggesting discrete-deconvolution methods might be necessary to detect touch-tuned neurons. To test this, in each of the 9 sets of time-series we identify touch-tuned neurons by a significant peak in their touch-triggered activity (Fig 6a). Figure 6b shows that, while all data-sets have touch-tuned neurons, the number of such neurons differs substantially between them. And rather than being essential to detecting fast responding touch-tuned neurons, discrete deconvolution methods disagree strongly on touch-tuning, with LZero (events) finding more touch-tuned neurons than in the raw calcium, but MLSpike (events) finding less than half that number.

**Figure 6:**
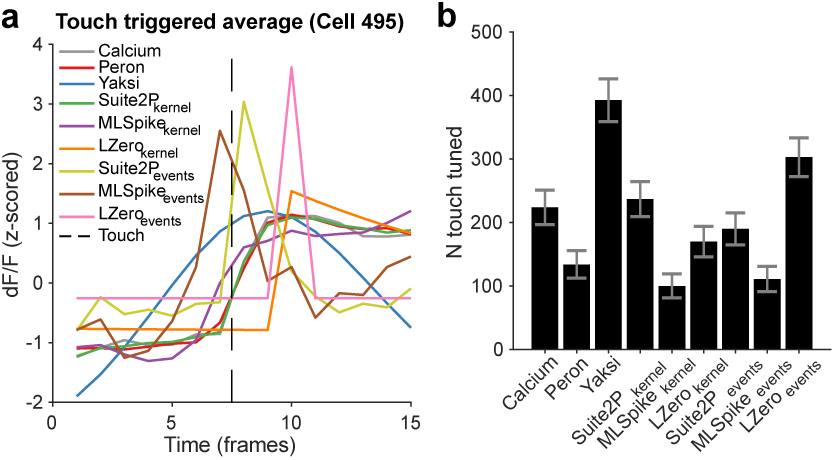
Touch-triggered neuron responses. (a) Touch-triggered average activity from one neuron, across all deconvolution methods. The dotted line is the imaging frame in which the whisker touched the pole. (b) Number of touch-tuned neurons across deconvolution methods. A neuron is classed as touch-tuned if its peak touch-triggered activity is significantly greater than randomly resampled data. Error bars are 95% Jeffreys confidence intervals for binomial data.

Thus our inferences of the coding of task-wide or specific sensory events crucially depends on both whether we deconvolve the raw calcium time-series or not, and on which algorithm we choose to do so.

### 2.6 Inconsistent recovery of population correlation structure across deconvolution approaches

The high yield of neurons from calcium imaging is ideal for studying the dynamics and coding of neural populations (Harvey et al., 2012; Huber et al., 2012; Kato et al., 2015). Many analyses of a population’s dynamics or coding start from pairwise correlations between its neurons, whether as measures of a population’s synchrony or joint activity, or as a basis for further analyses like clustering and dimension reduction (Cunningham and Yu, 2014; Humphries, 2017; Stringer et al., 2019b). Consequently, differences in correlation estimates will play out as different inferences of population dynamics or population coding. For example, weak correlations between neurons in primary visual cortex would be evidence of sparse coding of visual information (Stringer et al., 2019a; Rumyantsev et al., 2020). We now ask how our inferences of population correlation structure in the barrel cortex data also depend on the choice of deconvolution method.

Figure 7a shows that the distributions of pairwise correlations qualitatively differ between the sets of time-series we derived from the same calcium imaging data. The considerably narrower distributions from the discrete deconvolution time-series compared to the others is expected, as these time-series are sparse. Nonetheless, there are qualitative differences within the sets of discrete and continuous time-series. Some distributions are approximately symmetric, with broad tails; some asymmetric with narrow tails; the correlation distribution from the Peron method time-series is the only one with a median below zero. These qualitative differences are not due to noisy estimates of the pairwise correlations: for all our sets of time-series the correlations computed on a sub-set of time-points in the session agree well with the correlations computed on the whole session (Figure 7b). (Although we note that, as expected, the three spike-event time-series require far more time-points to obtain stable correlation estimates, because of their sparse events). Thus pairwise correlation estimates for each method are stable, but their distributions differ between methods.

**Figure 7:**
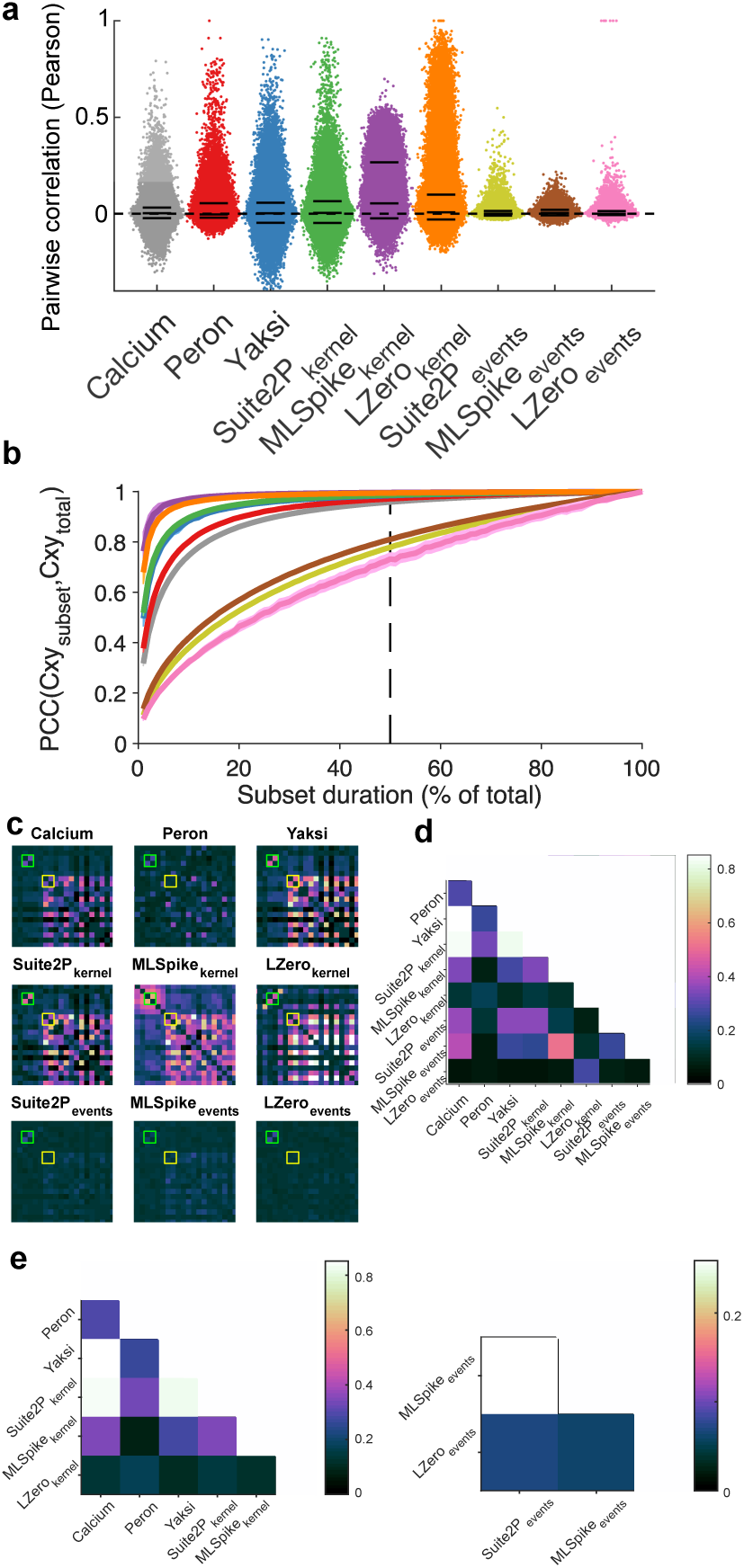
Effects of deconvolution on pairwise correlations between neurons. (a) Distributions of pairwise correlations between all neurons, for each deconvolution method (one dot per neuron pair, x-axis jitter added for clarity). Solid black lines are 5th, 50th and 95th percentiles. (b) Stability of correlation structure in the population. We quantify here the stability of the pairwise correlation estimates, by comparing the correlation matrix constructed on the full data (*Cxy*_*total*_) to the same matrix constructed on a subset of the data (*Cxy*_*subset*_). Each data-point is the mean correlation between *Cxy*_*total*_ and *Cxy*_*subset*_; one line per deconvolution method. Shaded error bars are one standard deviation of the mean across 100 random subsets. (c) Examples of qualitatively differing correlation structure across methods. Each panel plots the pairwise correlations for the same 50 neurons on the same colour scale. As examples, we highlight two pairs of neurons: one consistently correlated across different methods (green boxes); the other not (yellow boxes). (d) Comparison of pairwise correlation matrices between deconvolution methods. Each square is the Spearman’s rank correlation between the full-data correlation matrix for that pair of methods. We use rank correlation to compare the ordering of pairwise correlations, not their absolute values. (e) as in (d), but split to show continuous methods (left) or discrete deconvolution methods (right).

Looking in detail at the full correlation matrix shows that even for methods with similar distributions, their agreement on correlation structure is poor. Some neuron pairs that appear correlated from time-series processed by one deconvolution method are uncorrelated when processed with another method (Figure 7c). Over the whole population, the correlation structure obtained from the raw calcium, Yaksi and Suite2p (kernel) time-series all closely agree, but nothing else does (Figure 7d): the correlation structure obtained from LZero agrees with nothing else; and the discrete deconvolution methods all generate dissimilar correlation structures (Figure 7e). Our inferences about the extent and identity of correlations within the population will differ qualitatively depending on our choice of deconvolution method.

### 2.7 Deconvolution methods show the same population activity is both low and high dimensional

Dimensionality reduction techniques, like principal components analysis (PCA), allow researchers to make sense of large scale neuroscience data (Chapin and Nicolelis, 1999; Briggman et al., 2005; Churchland et al., 2012; Harvey et al., 2012; Cunningham and Yu, 2014; Kobak et al., 2016), by reducing the data from *N* neurons to *d < N* dimensions. Key to such analyses is the choice of *d* dimensions, a choice guided by how much of the original data we can capture. Differences in dimensionality imply different computations: for example, low-dimensional activity implies a sensory population uses a redundancy code, while high-dimensional activity implies the population uses a sparse code (Wohrer et al., 2013). To assess such inferences of population dimensionality, we apply PCA to our 9 sets of imaging time-series to estimate the dimensionality of the imaging data (which for PCA is the variance explained by each eigenvector of the data’s covariance matrix).

Figure 8a plots for each deconvolution method the cumulative variance explained when increasing the number of retained dimensions. Most deconvolution methods qualitatively disagree with the raw calcium data-set on the relationship between dimensions and variance. This relationship is also inconsistent across deconvolution methods; indeed the discrete deconvolution methods result in the shallowest (MLSpike_*events*_) and amongst the steepest (LZero_*events*_) relationships between increasing dimensions and variance explained. The number of dimensions required to explain 80% of the variance in the data ranges from *d* = 125 (Peron) to *d* = 1081 (MLSpike_*events*_), a jump from 8% to 70% of all possible dimensions (Fig 8b). Thus we could equally infer that the same L2/3 population activity is low dimensional (*<*10% dimensions required to explain 80% of the variance) or high-dimensional (*>*50% of dimensions required) depending on our choice of imaging time-series.

**Figure 8:**
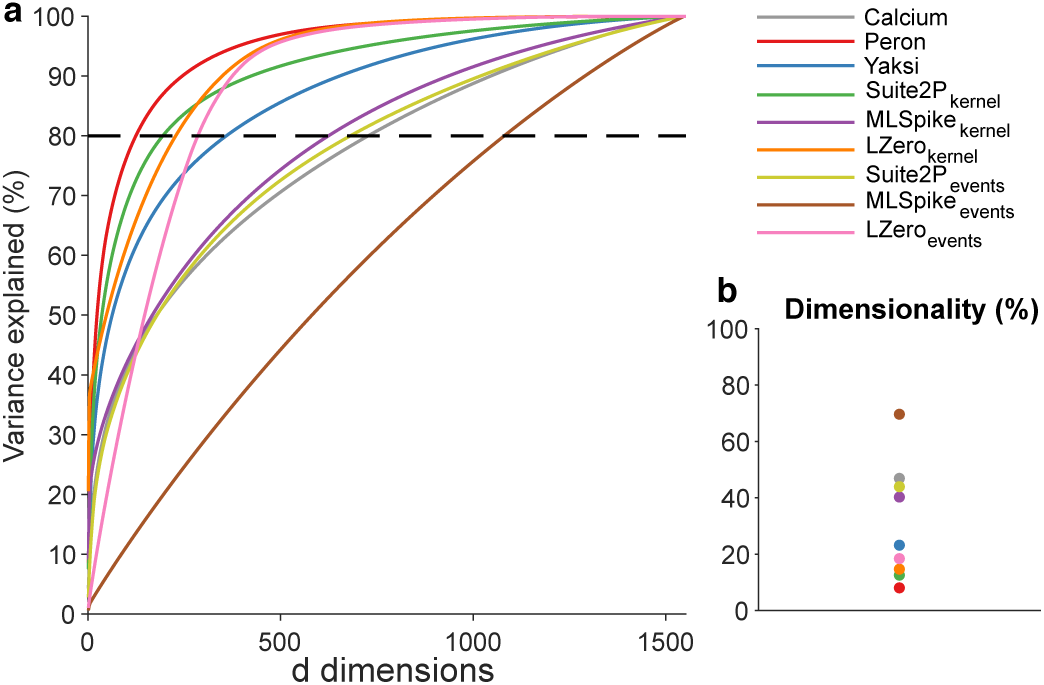
Dimensionality of population activity. (a) Cumulative variance explained by each dimension of the data’s covariance matrix, one line per deconvolution method. Dashed line is the 80% threshold used in panel (b). (b) Proportion of dimensions required to explain 80% of the variance in the data.

## 3 Discussion

Imaging of somatic calcium is a remarkable tool for capturing the simultaneous activity of hundreds to thousands of neurons. But the time-series of each neuron’s calcium fluorescence is inherently noisy and non-linearly related to its spiking. We sought here to address how our choice of corrections to these time-series – to use them raw, deconvolve them into continuous time-series, or deconvolve them into discrete events – affect the quality and reliability of the scientific inferences drawn. Our approach was to replicate the process of a typical population calcium-imaging study: choose an algorithm, choose its parameters using some reasonable heuristics, and analyse the resulting time-series.

Our results show the choice of processing qualitatively changes the potential scientific inferences we draw about the activity, coding, and correlation structure of a neural population in barrel cortex. Only the raw calcium and two of the processed time-series correctly capture the expected long-tailed distribution of spiking activity across the population. Neurons identified as being tuned to any feature of a pole-detection task differ widely between processing methods. Few methods agree on the pairwise correlation structure of the population. Moreover, the apparent dimensionality of the population activity can differ by an order of magnitude across the processing methods. Across all analyses, we consistently observe that the results differ sharply between the raw calcium and most, if not all, of the processed time-series. However, the deconvolved time-series also consistently disagreed with each other, even between methods of the same broad class (continuous or discrete time-series).

### 3.1 Accurate discrete deconvolution is possible, but sensitive

We find much that is encouraging. In fitting discrete deconvolution methods to ground-truth data, we found they can in principle accurately recover known spike-times from raw calcium time-series. A caveat here is that the choice of metric for evaluation and fitting of parameters is of critical importance. The widely-used Pearson correlation coefficient is a poor choice of metric as it returns inconsistent results with small changes in algorithm parameters, and leads to poor estimates of simple measures such as firing rate when used across methods and sampling rates. By contrast, the Error Rate metric (Deneux et al., 2016; Victor and Purpura, 1996) resulted in excellent recovery of ground-truth spike trains. Other recently developed methods for comparing spike-trains based on information theory (Theis et al., 2016) or fuzzy set theory (Reynolds et al., 2018), may also be appropriate.

However, while good estimates of ground-truth spike times can be achieved with modern discrete deconvolution methods (Berens et al., 2018; Pachitariu et al., 2018), the best parameters vary substantially between cells, and small changes in parameter values result in poor performance. This variation and sensitivity of parameters played out as widely-differing results between the three discrete deconvolution methods in analyses of neural activity, coding, and correlation structure.

### 3.2 Choosing parameters for deconvolution methods

A potential limitation of our study is that we use a single set of parameter values for each discrete deconvolution method applied to the population imaging data from barrel cortex. But then our situation is the same as that facing any data analyst: in the absence of ground-truth, how do we set the parameters? Our solution here was to use the modal parameter values from ground-truth fitting. We also felt these were a reasonable choice for the population imaging data from barrel cortex, given that the ground-truth recordings came from the same species (mouse) in the same layer (2/3) of a different bit of primary sensory cortex (V1). It would be instructive in future work to quantify the dependence of analyses of neural activity, coding, and correlation on varying the parameters of each deconvolution method.

### 3.3 Ways forward

How then to solve the problem of the wide disagreements we report here, both between the raw calcium and the deconvolved time-series, and between the outputs of the deconvolution methods?

The simplest approach is to side-step the issue, and just use the raw calcium time-series. Many studies use the raw calcium signal as the basis for all their analyses (Harvey et al., 2012; Huber et al., 2012; Chu et al., 2016), perhaps assuming this is the least biased approach. Our results suggest caution: the discrepancy between the raw and deconvolved calcium on single neuron coding suggests an extraordinary range of possible results, from about half of all neurons tuned to the task down to less 5 percent. The qualitative conclusion – there is coding – is not satisfactory. Moreover, as noted by (Sabatini, 2019), the raw calcium fluorescence signal is a low-pass filtered version of the underlying spike train, which places strong limits on the maximum correlation between the raw signal and underlying spikes, and hence on any correlations between the raw signal and the behavioural variables related to those spikes. Indeed, the desire for better recovery of the spikes and their correlations with behaviour is one of the principle reasons for developing deconvolution methods.

A natural step then is to improve deconvolution methods with better forward models, like MLSpike, for the link from spiking to calcium fluorescence (Greenberg et al., 2018). Indeed, as sensors with faster kinetics (though fundamentally limited by kinetics of calcium release itself) and higher signal-to-noise ratios are developed (Badura et al., 2014; Dana et al., 2016, 2019), so the accuracy and robustness of de-noising and deconvolution should improve; and as the neuron yield continues to increase (Ahrens et al., 2013; Stringer et al., 2019a), so the potential for insights from inferred spikes or spike-driven events grows. Developing further advanced deconvolution algorithms will harness these advances, but are potentially always limited by the lack of ground-truth to fit their parameters (Wei et al., 2019). Worse, no matter how good the forward model for a single neuron, our results suggest the wide variation in the model parameters needed for each neuron would make population analyses challenging to interpret.

A simple alternative approach to the inconsistencies between different forms of deconvolved time-series is to triangulate them, and take the consensus across their results. For example, our finding of a set of tuned neurons across multiple methods is strong evidence that neurons in L2/3 of barrel cortex are responsive across the stages of the decision task. Further examples of such triangulation in the literature are rare; Klaus and colleagues (Klaus et al., 2017) used two different pipelines to derive raw Δ*F/F* of individual neurons from one-photon fibre-optic recordings in the striatum, and replicated all analyses using the output of both pipelines. Our results encourage the further use of triangulation to create robust inference: obtaining the same result in the face of wide variation increases our belief in its reliability (Munafò and Davey Smith, 2018).

There are caveats to triangulating by using a full consensus across three or more versions of the time series. For single neuron analyses, such a full consensus inevitably comes at the price of reducing the yield of neurons to which we can confidently assign roles. There is also an assumption that all contributions to the consensus contain useful data: if one deconvolution method returns time-series with no relation to the underlying spike events, then including its outputs in the consensus would inevitably worsen the results. An alternative version of triangulation partially circumventing these problems would be to separately take the consensus between the raw calcium time-series and each of two or more spike-event deconvolution methods, and then combine the results. Future work on triangulation approaches would also need to look at how to combine more complex analyses than single neuron properties, such as pairwise correlations.

Another approach, little explored to date, would be to use data constraints to tune the deconvolution algorithm parameters. One option would be to use known properties of neural activity in a recorded population as constraints. We showed, for example, that some deconvolution methods did not recover the expected population-wide distribution of activity in layer 2/3 of barrel cortex; so constraining all algorithms to reproduce the longtailed activity distribution may improve agreement between them in measures of coding and correlation. Another option would be to tune deconvolution parameters to maximise consistency within the deconvolved data. For example, Pachitariu et al. (2018) recently proposed maximising the correlation between deconvolved traces from the same neuron obtained between trials of the same visual stimulus. Such an approach needs a suitable task design to ensure consistent conditions within which to compare responses of the same neuron (such as identical duration repeats of identical visual stimuli) – and which therefore could not be applied to the pole-detection task considered here. It would also require that the known variations in a neuron’s response between repeats of the same task condition or stimulus is not large enough to prevent meaningful correlations between repeats.

Our results provide impetus for different directions of research, not just to improving our modelling of the relationship between spikes and the somatic calcium signal, but also focussing on how we can verify results across the output of different deconvolution algorithms, and thus provide robust scientific inferences about neural populations.

## 4 Methods

### Ground truth data

Ground truth data were accessed from crcns.org (Svoboda, 2015), and the experiments have been described previously (Chen et al., 2013). Briefly, mouse visual cortical neurons expressing the fluorescent calcium reporter protein GCaMP6s were imaged with two-photon microscopy at 60Hz. Loose-seal cell-attached recordings were performed simultaneously at 10kHz. Recordings were made in awake mice during 5 trials (4s blank, 4s stimulus) of the optimal moving grating stimulus (1 of 8 directions) for the cell-attached neuron. The data-set contains twenty one recordings from nine neurons.

Neuropil subtraction was performed as described in Chen et al. (2013), based on example code provided alongside the data at crcns.org. Neuropil signals – defined as the average fluorescence from all pixels within a 20 *µ*m radius from each celcentre excluding the region of interest (ROI) – were subtracted from cell fluorescence in a weighted fashion, *F*_*corrected*_ = *F*_*cell*_ − 0.7*F*_*neuropil*_.

### Population imaging data description

Population imaging data was accessed from crcns.org and have been described previously (Peron et al., 2015b). Briefly, volumetric two photon calcium imaging of primary somatosensory cortex (S1) was performed in awake head-fixed mice performing a whisker-based object localisation task. In the task a metal pole was presented in one of two locations and mice were motivated with fluid reward to lick at one of two lick ports depending on the location of the pole following a brief delay. Two photon imaging of GCaMP6s expressing neurons in superficial S1 was performed at 7Hz. Images were motion corrected and aligned, before regions of interest were manually set and neuropil-subtracted. A single recording from this dataset was used for population analysis. The example session had 1552 neurons recorded for a total of 23559 frames (56 minutes).

### List of deconvolution methods

#### MLSpike

`MLSpike` (Deneux et al., 2016) was accessed from https://github.com/mlspike. MLSpike uses a model-based probabilistic approach to recover spike trains in calcium imaging data by taking baseline fluctuations and cellular properties into account. Briefly, MLSpike implements a model of measured calcium fluorescence as a combination of spike-induced transients, background (photonic) noise and drifting baseline fluctuations. A maximum likelihood approach determines the probability of the observed calcium at each time step given an inferred spike train generated through a particular set of model parameters. MLSpike returns a maximum a posteriori spike train (as used here), or a spike probability per time step.

MLSpike has a number of free parameters, of which we optimise three: *A*, the magnitude of fluorescence transients caused by a single spike; *tau*, calcium fluorescence decay time; *sigma*, background (photonic) noise level. MLSpike also has parameters for different calcium sensor kinetics (for OBG, GCaMP3, GCaMP6 and so on) which we fix to default values for GCaMP6.

For our analysis of event rate MLSpike’s spike train was counted (mean event count per second), and for subsequent analyses was converted to a dense array of spike counts per imaging frame.

#### Suite2P

`Suite2P` (Pachitariu et al., 2016, 2018) was accessed from https://github.com/cortex-lab/Suite2P. Suite2P was developed as a complete end-to-end processing pipeline for large scale 2-photon imaging analysis - from image registration to spike extraction and visualization - of which we only use the spike extraction step. The spike deconvolution of Suite2P uses a sparse non-negative deconvolution algorithm, greedily identifying and removing calcium transients to minimise the cost function

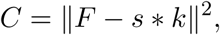

where the cost *C* is the squared norm of fluorescence *F* minus a reconstruction of that signal comprising a sparse array of spiking events *s* multiplied by a parameterised calcium kernel *k*. The kernel was parameterised following defaults for GCaMP6s (exponential decay of 2 seconds, though it has been shown the precise value of this parameter does not affect performance for this method (Pachitariu et al., 2018)).

Suite2P has a further free parameter which sets the minimum spike event size, the *Threshold*, which determines the stopping criteria for the algorithm.

Elements of *s* are of varying amplitude corresponding to the amplitude of the calcium transients at that time. For ground truth firing rate analysis we are interested in each algorithm’s ability to recover spike trains, therefore we treat each event as a ‘spike’ and optimise the algorithm appropriately. For our analysis of event rate Suite2P’s event train was counted (mean event count per second), and for subsequent analyses was converted to a dense array of varying amplitude events (i.e. *s*) per imaging frame.

#### LZero

The method we refer to as `LZero` was written in Matlab based on an implementation in *R* accessed at https://github.com/jewellsean/LZeroSpikeInference. A full description is available in the paper of Jewell and Witten (2018). Briefly, in LZero spike detection is cast as a change-point detection problem, which could be solved with an *l*_0_ optimization algorithm. Working backwards from the last time point the algorithm finds time points where the calcium dynamics abruptly change from a smooth exponential rise. These change points correspond to spike event times. Spike inference accuracy is assessed similarly to Suite2P by measuring the fit between observed fluorescence and a reconstruction based on inferred spike times and a fixed calcium kernel.

LZero has two free parameters - *lambda*, a tuning parameter that controls the trade-off between the sparsity of the estimated spike event train and the fit of the estimated calcium to the observed fluorescence; and *scale*, the magnitude of a single spike induced change in fluorescence.

For our analysis of event rate LZero’s spike train was counted (mean event count per second), and for subsequent analyses was converted to a dense array of spikes per imaging frame (maximum one spike per imaging frame due to limitations of the algorithm).

#### Yaksi

`Yaksi` is an implementation of the deconvolution approach of Yaksi and Friedrich (2006). The fluorescence time series is low-pass filtered (4^th^ order butterworth filter, 0.7Hz cutoff) to remove noise before having a calcium kernel (exponential decay of 2 seconds, as used in Suite2P and LZero above) linearly deconvolved out of the signal using Matlab’s `deconv` function. The output of Yaksi is a continuous signal approximating spike density per unit time.

#### Peron events

`Peron events` refer to the de-noised calcium event traces detailed in the original Peron et al. (2015b) paper. Here a version of the ‘peeling’ algorithm (Lütcke et al., 2013) was developed, a template-fitting algorithm with variable decay time constants across events and neurons. The output for analysis is a continuous signal approximating de-noised calcium concentration per unit time.

#### *Events* and *kernel* versions of spike inference methods

Where a spike inference method returns spike counts per time point, these are plotted as Method_*events*_. To compare to other methods that return a de-noised dF/F or firing rate estimates, these event traces are convolved with a calcium kernel and plotted as Method_*kernel*_. The kernel used is consistent with that used as a default for GCaMP6s in MLSpike, Suite2P and LZero, namely an exponential decay of two seconds duration normalised to have an integral of 1.

### Ground truth spike train metrics

Pearson correlation coefficient was computed between the ground truth and inferred spikes (MLSpike) or events (Suite2P, LZero) following convolution of both with a gaussian kernel (61 samples wide, 1.02 seconds).

Error Rate was computed between the ground truth and inferred spikes/events using the Deneux et al. (2016) implementation of normalised error rate, derived from Victor and Purpura (1996) – code available https://github.com/MLspike. Briefly, the error rate is 1 - F1-score, where the F1-score is the harmonic mean of sensitivity and precision (Davis and Goadrich, 2006),

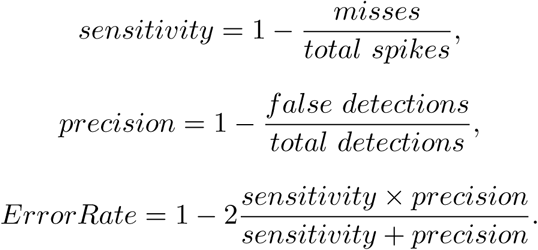

Hits, misses and false detections were counted with a temporal precision of 0.5 seconds. For normalised estimation of errors in firing/event rate we compute,

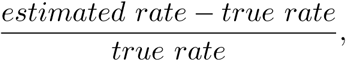

where spike/event rates are measured in Hz.

### Parameter fitting

For each method the best parameters for each neuron were determined by brute force search over an appropriate range (i.e. at least two orders of magnitude encompassing full parameter ranges used in the original publications for each method). The parameter ranges were explored on a log scale as follows: MLSpike A (0.01:1, 21 values), tau (0.01:5, 21 values), sigma (0.01:1, 21 values); Suite2P Threshold (0.1:100, 13 values); LZero lambda (0.1:20, 23 values), scale (0.1:20, 23 values).

The modal best parameters, as determined using Error Rate on downsampled data, were then fixed for the population imaging data analysis. These were: MLSpike A: 0.1995, tau: 1.9686, sigma: 0.0398; Suite2P Threshold: 1.7783; LZero sigma: 0.1; lambda: 3.1623.

### Downsampling

Ground truth calcium data was downsampled from 60Hz to 7Hz in Matlab by up-sampling by 7 (interpolating the signal) and then downsampling the resultant 420Hz time-series of frames to 7 Hz by sampling every 60th frame.

### 4.1 Event rate estimation

Spike inference methods (Suite2P_*events*_,MLSpike_*events*_, LZero_*events*_) return estimated spike times (MLSpike), or event times (Suite2P/LZero) which were converted into mean event rates (Hz) per neuron.

The event rate for continuous methods (Calcium, Peron, Yaksi, Suite2P_*kernel*_, MLSpike_*kernel*_, LZero_*kernel*_) for each neuron was determined by counting activity/fluorescence transients greater than three standard deviations of the background noise. Background noise was calculated by subtracting a four-frame moving average of the fluorescence from the raw data to result in a ‘noise only’ trace. This operation was done separately for each neuron and each method. Event rate was then computed in Hz.

Silent neurons were defined as neurons with event rates below 0.0083Hz (or fewer than one spike per two minutes of recording) as in O’Connor et al. (2010).

### 4.2 Task-tuned neurons

Task-tuning was determined for each neuron using the model-free approach of Peron et al. (2015b). Neurons were classed as task-tuned if their peak trial-average activity exceeded the 95th percentile of a distribution of trial-average peaks from shuffled data (10000 shuffles of time-series order). The shuffle test was done separately for correct lick-left and lick-right trials and neurons satisfying the tuning criteria in either case were counted as task-tuned.

Tuned neuron agreement was calculated as the number of methods that agreed to the tuning status of a given neuron, for all methods and separately for continuous and spike inference methods.

### 4.3 Touch-tuned responses

Touch-tuned neurons were determined by first computing touch-triggered average activity for each neuron, then calculating whether the data distribution of peak touch-induced activity exceeds the expected activity of resampled data. In more detail, the time of first touch between the mouse’s whisker and the metal pole on each trial was recorded. For each neuron, one second of activity (seven data samples) was extracted before and after the frame closest to the first touch of each trial (15 frames total per trial); taking the mean touch-triggered activity over trials gave the average touch response for the neuron. To determine whether the neuron was touch tuned or not, we compared the neuron’s peak mean response *r*_*data*_ to a null distribution by taking a randomly sampled 15 frame segment of a trial, finding the peak mean response across trials *r*_*null*_, and repeating this calculation for 10000 random samples. A p-value for the data peak response was calculated as *p* = #*{r*_*null*_ < *r*_*data*_*}/*10000. Over all neurons, a neuron was considered touch-tuned if *p* < 0.05 after Benjamini-Hochberg correction.

### 4.4 Pairwise correlations

Pairwise correlations (Pearson correlation coefficients, Fig. 7a) were calculated between all pairs of neurons at the data sampling rate (7Hz).

Stability of correlation estimates (Fig. 7b) at the recording durations used was assessed by computed the similarity between correlation distributions for the the intact dataset to those from subsets of the dataset. For each deconvolution method, we computed the pairwise correlation matrix using the entire session’s data, as above. We also sampled a subset of time-points (1%-100%) of the full dataset at random without replacement and computed a matrix of pairwise correlations for this subset. We then compute the similarity between the total and subset matrices using Pearsons correlation coefficient. This process was repeated 100 times and the mean (line) and standard deviation (shading) of the 100 repeats were plotted.

### 4.5 Correlations between correlation matrices

Correlations between correlation matrices (Fig. 7c-e) were computed using Spearman’s rank correlation between the unique pairwise correlations from each method (i.e. the upper triangular entries of the correlation matrix).

### 4.6 Dimensionality

To determine the dimensionality of each dataset we performed eigendecomposition of the covariance matrix of each dataset. The resultant eigenvalues were sorted into descending order *λ*_1_ ≥ *λ*_2_ ≥ …*λ*_*N*_, and the cumulative variance explained by *d* dimensions computed as 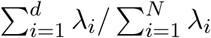.

## Notes

### Competing Interest Statement

The authors have declared no competing interest.

### Summary of Updates

Added full results for sensitivity analysis of third deconvolution algorithm to Figures 1 & 2. Added example of the variation of an inferred spike train across plausible parameters applied to a single neuron Clarified text throughout

